# Detecting T cell receptor rearrangements *in silico* from non-targeted DNA-sequencing (WGS/WES)

**DOI:** 10.1101/201947

**Authors:** Lara Lewis McGrath, Tristan J. Lubinski, J. Carl Barrett, Humphrey Gardner

## Abstract

To better understand the composition of heterogeneous tissue samples used in generating large genomic datasets, we developed a method for estimating the abundance of T cells within the cellular population. Somatic recombination of chromosomal DNA in T cells creates a vast repertoire of structurally divergent T cell receptors (TCRs) that recognize an array of non-self proteins. It also generates a genomic signature by which TCR sequences can be distinguished from other cell types in non-targeted NGS genomic data. Here we leverage this signature to extract reads with rearranged TCR sequences from a non-targeted population, such as whole genome sequencing (WGS) or whole exome sequencing (WES) datasets. We isolate and confirm T cell rearranged reads from the remainder of the genome (99.9%), accurately estimate relative T cell abundance within a cellular population, and provide a snapshot of the T cell receptor repertoire. This approach is unique from available TCR software options that focus on examining the overall diversity of the TCR repertoire and require prior amplification or selection of this region before sequencing, and has particular utility in immunoscoring clinical patient samples in situations where genomic data exists and other approaches are unavailable.

## 1. Introduction

T cell receptors (αβ or γδ) are dimeric protein complexes formed by two chains derived from separate somatic recombination events. During recombination in the α and γ loci, the variable (V) and joining (J) gene segments combine into an intact VJ region. A similar process occurs in β and δ recombination, with inclusion of one or more nucleotides from the short diversity (D) gene segment flanked by random nucleotides on either side, creating a hypervariable CDR3 (complementary determining region 3) within the intact V(D)J region (Davis and Bjorkman, 1988). The CDR3 is the principal site of contact for recognition specificity with foreign antigens presented by the major histocompatibility complex (MHC), and provides the ability to recognize a diverse set of foreign peptides. The observable diversity of CDR3 sequences is often generated as useful measure of the TCR repertoire, or diversity of αβ and γδ chains, as αβ T cells account for 95-99% of T cells. Statistical inference of this diversity estimates 10^14^ distinct CDR3 clonotypes (Murugan, et al., 2012), though the realized repertoire is estimated in the range of 1-4 × 10^6^ for healthy individuals (Robins, et al., 2009; Robins, et al., 2010; Warren, et al., 2011).

At present, several software applications exist for annotating T cell repertoire from NGS data, including IgBlast (NCBI) (Ye, et al., 2013), IMGT/High-V Quest (Alamyar, et al., 2012), MiXCR (Bolotin, et al., 2015), MIGEC (Shugay, et al., 2014), and IgRepertoireConstructor (Safonova, et al., 2015). These approaches are becoming increasingly sophisticated and accurate, yet they share the prerequisite that sequencing data has been selected for the VDJ recombination region during library construction, either through custom primers or other targeted methods. Rather than a measure of the repertoire of an individual, for which these targeted approaches are the gold standard, our approach aims to estimate αβ T cell abundance within the cellular population by detecting VDJ rearrangements *in silico* from NGS data without prior amplification.

## 2. Method

Extracting reads containing VDJ rearrangements from an unselected population necessitates bioinformatically detecting reads inherently diverse in nature and accurately reducing false positive calls from off target reads. We overcome these challenges by employing a step-wise method to isolate and confirm TCR β rearrangements using read files (fastq) from DNA-sequencing:

Align - rapidly isolate reads with homology to both V and J genes (kmer based seed-and-score alignment algorithm from MiXCR (Bolotin, et al., 2015))
Filter - subset potential VDJ rearrangements from aligned reads by filtering based on prior knowledge
Validate - confirm true VDJ rearranged reads with sensitive BLAST alignments

Through this workflow, our method (TCRbiter) acts as an arbiter for distinguishing TCR rearrangements from reads spanning the remainder of the genome (99.9+%) (Figure 1A). And by normalizing the number of detected VDJ rearranged reads against read coverage in this region, we are able to predict relative T cell abundance within a population.

**Figure 1A.**
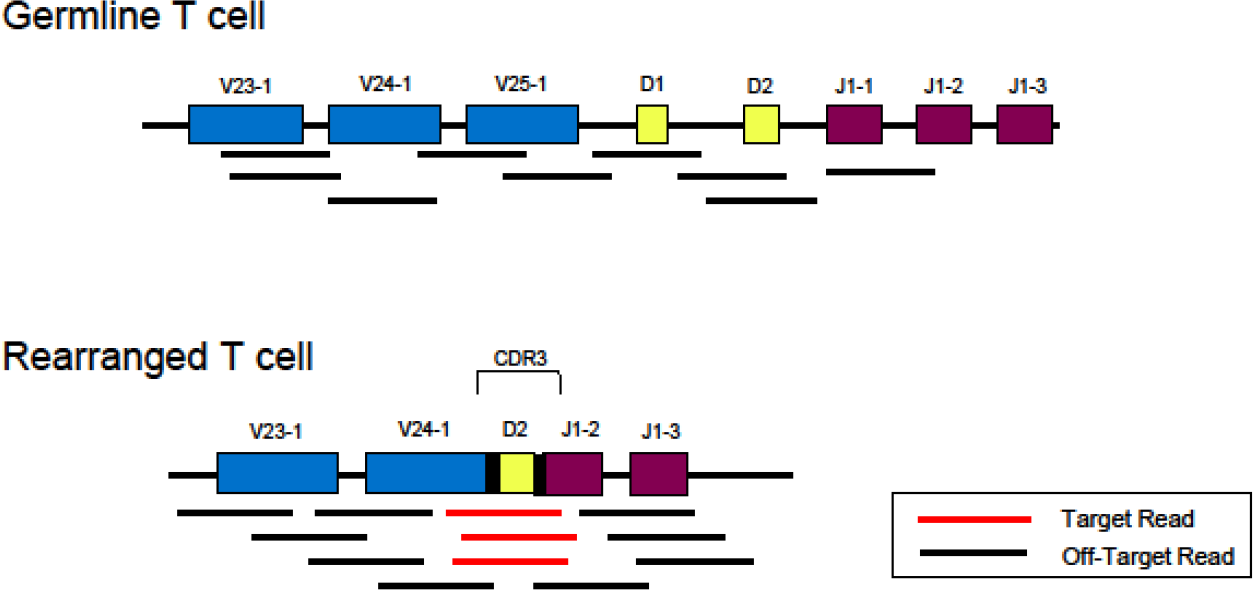
Schematic representation of target and off-target reads for detecting T cell rearrangements from genomic DNA sequences. Target reads for our method span the hypervariable CDR3 and include both V and J gene segments. Off target reads from this region may include either a V, D, or J gene segment and/or intronic sequences. Off target reads could be from a germline T cell or any cell type other than a rearranged T cell, and also outside of this region (not depicted). Not drawn to scale. For reference, average germline gene segment lengths are 471 bp (V), 13 bp (D) or 49 bp (J).

We developed and refined this approach using real and *in silico* generated true and false positive data sets. We tested our method against *in silico* simulations and real clinical data, and benchmarked it against comparable software and methods. Additional details on the methodology and validation steps are provided in the supplementary materials (Figures S1-4). Our method is open source and freely available to use as a Python program running on all Unix-compatible platforms. The source code is available from github.com/AstraZeneca-NGS/tcrbiter

## 3. Results

We offer this bioinformatic pipeline as a method for immunoscoring genome or exome sequencing data, finding particularly utility in situations where sample quantity is limited and more targeted approaches are unfeasible or unavailable. To demonstrate the ability of this workflow to produce relative T cell abundance in such a setting, we employed our technique against whole exome sequencing (WES) datasets from The Cancer Genome Atlas (TCGA) and compared it to determination of T cell infiltration performed by histological review.

### Comparison of matched tumor pairs by WES and histology

Using a subset of Thymoma (THYM) primary tumor samples from TCGA, we interrogated WES reads with matched tumor slides. A pathologist (H.G.) quantified lymphocyte counts by slide review (Figure 1B). We identified 0-19 VDJ rearranged reads per sample and share this value as a percentage (% TCR Reads) of the total coverage from this region for each sample, which was on average 42x coverage. Percent TCR reads detected were significantly (*p* < 0.05) correlated to T cell infiltrate levels as determined by pathology (*F*_1,8_ = 6.68; *p* value = 0.032).

**Figure 1B.**
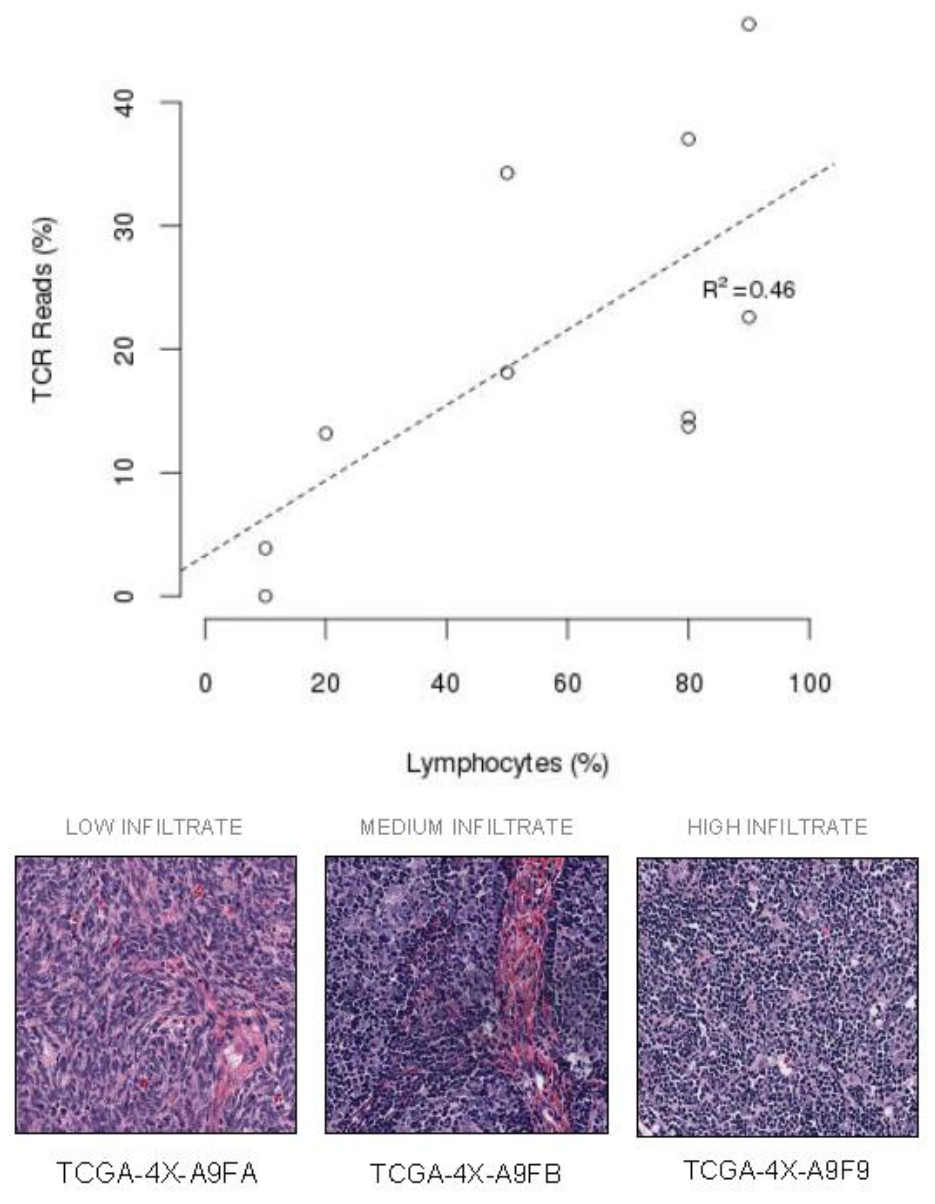
Detection of T cell infiltrate levels in WES compared to lymphocyte counts by examination of pathology on ten matched TCGA Thymoma tumor samples. TCR Reads (%) indicates the number of detected reads with T cell rearrangements normalized by the average coverage depth for the TCR loci. Slides displayed represent individual samples included in this set that are representative of low, medium,and high lymphocyte infiltration. Relative T cell infiltration (% TCR Reads) detected was significantly related (*p* < 0.05) to percent lymphocyte infiltration determined by pathology.

Moreover, from this analysis we detected a combined 81 TCR reads across all ten surveyed patients, which we re-annotated using IgBlast (Ye, et al., 2013). These 81 reads annotated to 63 unique clonotypes, specified by common CDR3 amino acid sequence. 71.43% (45/63) of clonotypes were productive and, of these, 6.67% (3/45) were shared, meaning they were identified in tumor samples from two or more different patients.

## 4. Conclusions

Here we present and validate a novel method to repurpose genomic data to provide a relative T cell infiltration level, thus combining a genomic and immune evaluation from a single sample portion.

In this study, we show our method is highly effective at discriminating between T cell rearranged reads and germline reads (Figure S2,S4). We applied this approach to examples from a clinical setting, examining WES and histological assays from primary tumor samples in TCGA. Results indicate the percent of TCR reads (as a fraction of total coverage) is significantly correlated to lymphocyte counts determined by histology (*p* < 0.05).

Recent studies employed related approaches to assemble consensus sequences for VDJ rearrangements from bulk tissue RNA-Seq (Brown, et al., 2015; Li, et al., 2016) and DNA data (Levy, et al., 2016). In comparing DNA- vs RNA-derived methodologies, the advantage of working with RNA is a higher proportion of detectable reads. However, by using DNA-derived data, we eliminate the possibility of multiple mRNA transcripts with the same sequence originating from a single cell, rather than multiple cells within an expanded clonal population. This distinction allows for more accurate quantification of clonal frequency. Though at present, whether from transcriptome-wide gene expression studies or WES/WGS, sequencing depth presents an important limitation to our ability to detect a substantial portion of the clonal population. However, even without deep sequencing the TCR reads detected from our subset of thymoma primary tumors were sufficient in revealing 3 productive clonotypes shared between two or more patients. Across a greater number of samples, one can generate candidates for public clones with antigen-specificity in a particular indication and may have applications in adoptive T cell therapies.

In summary, interrogating DNA-sequencing data for shared clones and relative T cell infiltration levels supplies two valuable pieces of information that were previously undeterminable from WGS or WES assays. With clinical trial samples, sample procurement is finite and quantity is often limited. Experimentation on these samples necessitates the use of multiplexed assays that derive the most information and value. As the field expands in its investigation of both targeted therapies and immune-oncology approaches, this method can use DNA-sequencing data generated to understand the mutational landscape of our patients to also stratify samples by relative T cell infiltration. Moreover, in the future we can expect decreases in sequencing costs and improvements in complexity and coverage to allow for this method to capture greater proportions of the T cell repertoire and be meaningful in also assessing T cell diversity and clonal expansion.

## Acknowledgements

We are thankful to Jonathan Dry for his insight and expertise, and the name for our method: ‘TCRbiter’. We are also grateful to Elizabeth Maloney and Rory Kirchner for their knowledge and help.

